# Host-associated microbial diversity in New Zealand cicadas uncovers elevational structure and replacement of obligate bacterial endosymbionts by *Ophiocordyceps* fungal pathogens

**DOI:** 10.1101/2021.08.24.457591

**Authors:** Diler Haji, Jason Vailionis, Mark Stukel, Eric Gordon, Emily Moriarty Lemmon, Alan R. Lemmon, John P. McCutcheon, Chris Simon

**Affiliations:** Department of Integrative Biology, University of California, Berkeley, CA, USA; Department of Ecology & Evolutionary Biology, University of Connecticut, Storrs, CT, USA; Department of Biological Science, Florida State University, Tallahassee, FL, USA; Department of Scientific Computing, Florida State University, Tallahassee, FL, USA; Division of Biological Sciences, University of Montana, Missoula, MT, USA; Biodesign Center for Mechanisms of Evolution, Arizona State University, Tempe, AZ, USA

## Abstract

Host-microbe interactions influence eukaryotic evolution, particularly in the sap-sucking insects that often rely on obligate microbial symbionts to provision deficient nutrients in their diets. Cicadas (Hemiptera: Auchenorrhyncha: Cicadidae) specialize on xylem fluid and derive many essential amino acids and vitamins from intracellular bacteria or fungi (*Hodgkinia*, *Sulcia*, and *Ophiocordyceps*) that are propagated via transmission from mothers to offspring. Despite the beneficial role of these symbionts in nutrient provisioning, they are generally not considered to function within the gut where microbiota may play an important dietary role during insect diversification. Here, we investigate the relative impact of host phylogeny and ecology on gut microbial diversity in cicadas by sequencing 16S ribosomal RNA gene amplicons from 197 wild-collected cicadas and new mitochondrial genomes across 38 New Zealand cicada species, including natural hybrids between one species pair. We find a lack of phylogenetic structure and hybrid effects but a significant role of elevation in explaining variation in gut microbiota. Additionally, we provide evidence of *Hodgkinia* loss with gains of *Ophiocordyceps* fungal pathogens in all New Zealand cicadas examined that suggests convergent domestications of fungal pathogens. This highlights the macroevolutionary instability of obligate symbiosis and the relative importance of ecology rather than phylogeny for structuring gut microbial diversity in cicadas.

**Importance:** An unresolved question in evolutionary biology is how beneficial associations between eukaryotes and microbes impact macroevolutionary patterns. We report substantial data from natural populations that suggest the absence of macroevolutionary impacts from gut microbiota in cicadas. Instead, gut microbial diversity is better explained by elevational variation across an island landscape. Cicadas, like many insects, have obligate nutritional associations with bacteria housed in organs outside of the gut, but we show that these associations seem also to be unstable at macroevolutionary scales. We report evidence for unexpected and widespread replacement of obligate bacteria by a domesticated and formerly pathogenic *Ophiocordyceps* fungus representing an evolutionarily convergent pattern across the cicada phylogeny.

## Introduction

The study of patterns of associations between microbes and their hosts have provided many evolutionary insights (1), from the endosymbiotic origins of the eukaryotes (2) to the microbial basis of adaptations that are not directly encoded by the eukaryotic genome itself (3, 4). The reliance of hosts on functions provided by their microbes can lead to selection for specificity in host-microbe relationships when microbes are inherited through vertical transmission from mothers to offspring (5) or selected by hosts from the environment every generation (6). Often, such microbes are required for host development and are spatially segregated into specific cells and tissues within the body of a host individual (7-12).

Cicadas (Hemiptera: Auchenorrhyncha: Cicadidae) represent a well-studied example of obligate and specific associations between animals and microbes. They obtain essential amino acids and vitamins absent in their diets from two obligate symbionts, “*Candidatus* Sulcia muelleri” (hereafter *Sulcia*) and “*Candidatus* Hodgkinia cicadicola” (hereafter *Hodgkinia*). These microbes have largely non-overlapping metabolic functions required by their hosts for development and are housed in bacteriomes outside of the gut but within the cicada abdomen (12-16). While *Sulcia* has a more ancient association with insects in the suborder Auchenorrhyncha (17)*, Hodgkinia* is found specifically in the Cicadoidea and undergoes relatively rapid rates of mutation and genome fragmentation within host lineages unseen in other symbiont host lineages. It has been proposed that *Hodgkinia* genome fragmentation is due to the unusually long life-cycles of cicadas and high mutation rates in *Hodgkinia* compared to its co-symbiont *Sulcia*, which shows no genome fragmentation (11, 16, 18–20). As previously predicted by early microscopy (21, 22), recent molecular work shows that the loss of *Hodgkinia* is coincident with replacements by yeast-like *Ophiocordyceps* fungi (23). Remarkably, these yeast endosymbionts appear to be domesticated from cicada pathogens and provide the same nutrients as *Hodgkinia* (23). Similarly, yeast-like symbionts have been found to replace *Sulcia* rather than its co-partner endosymbiont in other Hemipteran insects, e.g., Fulgoroidea (24, 25).

Although these obligate symbionts – *Sulcia, Hodgkinia*, and *Ophiocordyceps* – are important in supplying nutrients to cicada hosts, they are not known to reside in the cicada gut. Comparatively little is known about cicada gut-associated microbes (26-28) despite the increasingly recognized nutritional role of animal gut microbiota (8, 29–31). As opposed to the specificity of obligate symbionts, gut microbiota may be facultative or transient with little specificity and involve complex microbial communities with varying functions across fluctuating ecological conditions (32), but may nonetheless be essential for host development (33-36).

Variation in microbial communities among animal hosts may be explained by many factors, including the host phylogeny itself (31, 37, 38), particularly when microbes are inherited. However, many ecological correlates have also been shown to impact host-associated microbial communities, particularly the conditions of the gut niche which experiences continuous input of environmental microbes (39-41). It remains unclear, however, to what extent microbial community assembly within many host species is influenced by fluctuating ecological conditions as compared to host evolution. Indeed, transient microbes may not have functions relevant to host fitness (42, 43). Yet these microbes may be a source of adaptive functions of novel microbial metabolism as hosts diversify (44). Broader surveys are needed to enhance our understanding of the causes and consequences of variation in host-associated gut microbiota and whether they play important roles in organisms with other symbiotic associations such as cicadas.

In this study, we generate 16S rRNA gene amplicons of microbial communities and newly assembled mitochondrial genomes across 38 New Zealand (NZ) cicada species and hybrids between one species pair to understand the relative contributions of host phylogeny and ecology to variation in gut microbiota. These 38 species comprise three genera and represent a radiation of species from a single colonizing ancestor which now occupy a wide variety of habitats and elevations. We show that phylogenetic distances among cicada hosts are poor predictors of gut microbial communities compared to elevational differences, suggesting little host specificity and a prominent role for ecological heterogeneity. Further supporting this lack of host specificity, we found no evidence of significant differences in gut microbial communities between hybrid and parental individuals. Serendipitously, our characterization of microbial communities produced many non-target 16S rRNA gene amplicons that provide evidence across all NZ cicadas of what is likely the absence of *Hodgkinia* symbionts and the widespread presence of abundant *Ophiocordyceps* fungal symbionts. This suggests convergent domestication of fungal pathogens in NZ cicadas and extends the findings of (23) for cicadas with a present-day distribution in Japan to the southern hemisphere. Our findings suggest that the interaction between cicadas and microbes at macroevolutionary time scales are strongly driven by turnover in the identity of obligate symbionts, in this case from bacterial to fungal symbionts, but gut microbiota are more strongly influenced by ecological variation than by host evolutionary history.

## Methods

### Sample collection and processing

#### Collection and outgroup choice

We collected New Zealand cicadas in 95% ETOH from 1995 to 2018 and stored them at -80 Celsius until processing (See Table S1 for specimen details). Based on previous work (45), we included several *Caledopsalta* (Cicadettini, Cicadettinae) cicada specimens from New Caledonia as an outgroup comparison to the New Zealand cicada clade comprising *Kikihia, Maoricicada*, and *Rhodopsalta.* We also included several *Magicicada* (Lamotialnini, Cicadettinae) from North America to represent members of the same subfamily with known presence of both *Hodgkinia* and *Sulcia* bacterial symbionts. Although we focus on the aforementioned groups, our dataset also includes, for comparative purposes, *Neotibicen* (North American, Cryptotympanini, Cicadinae) and *Platypedia* (North America, Tibicininae) specimens (Fig. S1-S3) but these are not discussed as part of our main findings. In summary, we caught cicadas by net or lured them using manually produced mating clicks, leveraging the attraction that male cicadas have for wing flicking sounds and movements associated with conspecific females (46, 47). We identified each specimen to species in the field using a combination of song, morphology, and knowledge of their evolutionary history and distribution (48-50). One hybrid-descended lineage, *K.* “muta x tuta”, was sampled. This lineage possesses *K. muta* nuclear DNA, song, and morphology, but *K.* “tuta” mtDNA as a result of ancient hybridization and introgression of the mitochondrion (50). The identity of *K.* “muta x tuta’’ samples were inferred based on the geographic distribution of the *K. muta* and *K.* “muta x tuta’’ lineages. For specimens sampled from localities known to have both lineages, we sequenced the mitochondrial COI gene and treated 100% matches to *K.* “tuta” sequences in GenBank as confirmation of *K.* “muta x tuta” identity.

#### Dissection

We sterilized specimens by submerging them in 2% household bleach, letting them sit for 1-2 minutes and then washing them in both 50-70% alcohol and sterile water. We then dissected specimens using small scissors, forceps and pins to access gut tissue ventrally. The complex structure of the cicada gut required prolonged (15-30 min) and relatively tedious dissection compared to insects with relatively simpler guts (Fig. S1). We either extracted both gut and reproductive tissue or only gut tissue depending on the specific dataset produced (dataset specifications provided below). Tissue samples were either directly placed into Powersoil bead tubes or stored in sterile cryotubes and kept frozen until DNA extraction. All dissection equipment was sterilized with 10% bleach and then treated with UV light in a crosslinker for at least one minute prior to dissection. We carried out dissections over the course of several months, with 2-15 dissections on any given day.

We binned processed samples into three sample batches based on the timing and methodological differences in processing, the workers who processed them, and sampling design: B1, B2, B3 (Table S1). Dataset-specific variations are described as follows:

#### B1 Dataset

Combined gut and male reproductive tissue from *Kikihia muta* and *Kikihia* “tuta” representing nine parental populations and six previously identified hybrid populations extracted using a Powersoil DNA Isolation kit (MO BIO Laboratories) under the standard protocol. The entire purification process was performed using the Powersoil DNA Isolation protocol (not DNeasy).

#### B2 Dataset

Gut tissue from a broad sampling of New Zealand cicada species mostly within *Kikihia* extracted by mechanical lysing within Powersoil bead tubes containing Powersoil lysis buffer and subsequent DNeasy 96-well plate extraction under standard protocols (beginning after Proteinase-K treatment and incubation).

#### B3 Dataset

Gut tissue from a broad sampling of New Zealand cicada species including outgroup species from New Caledonia and various North American cicadas extracted by mechanical lysing within Powersoil bead tubes containing Powersoil lysis buffer and subsequent DNeasy 96-well plate extraction under standard protocols (beginning after Proteinase-K treatment and incubation).

### Amplicon sequencing of 16s V4 rRNA

We amplified the V4 region of bacterial 16S rRNA using universal barcoded primers 515F (5’-GTGCCAGCMGCCGCGGTAA-3’) and 806R (5’-GGACTACHVGGGTWTCTAAT-3’) with attached Illumina-compatible adapters and indices (Microbial Analysis, Resources, and Services Facility, University of Connecticut) under the following PCR conditions: 95°C for 2 min, 35 cycles of 95°C for 30 sec, 55°C for 1 min, 72°C for 1 min, and then a final extension at 72°C for 5 min. All libraries were quantified with QIAxcel and manually inspected for proper marker alignment. We normalized libraries by pooling to the lowest concentration for each of the sample sets. Samples that could not be quantified due to low concentration were pooled using the maximum product available. Pooled samples were cleaned and size-selected using a bead-based approach. Final 16S amplicon libraries were sequenced paired end on Illumina MiSeq.

Dataset-specific variations are described as follows:

#### B1 Dataset

Amplified with V4 16S rRNA primers as above using EmeraldAmp GT PCR Master Mix (TAKARA BIO).

#### B2 Dataset

Amplified in a separate facility (Microbial Analysis, Resources, and Services Facility, University of Connecticut) with V4 16S rRNA primers as above using GoTaq DNA Polymerase (PROMEGA).

#### B3 Dataset

First amplified with primers 27F (5’-AGAGTTTGATCMTGGCTCAG-3’) and 1492R (5’-GGTTACCTTGTTACGACTT-3’) using EmeraldAmp GT PCR Master Mix (TAKARA BIO) to minimize the amplification of non-bacterial taxa under the following cycling conditions: 94°C for 5 min, then 5 cycles of 94°C for 45 sec, 56°C for 45 sec, 72°C for 1.5 min, and then a final extension at 72°C for 10 min. The product was used as a template for amplification with V4 16S rRNA primers as in dataset B1.

### Negative Controls

We took various controls when processing the B2 and B3 datasets, most of which were taken as part of the B3 dataset. Six PCR controls across the two datasets were taken (two controls in the B2 dataset and four controls in B3 dataset) during library amplification using V4 16s rRNA primers (Fig. S2, pcr and control). The remaining controls are associated with the B3 dataset and are as follows: Six dissection reagent controls of fluid used prior to dissection (Fig. S2, dissection), 10 controls of surface contents of forceps after transferring tissue from dissection plates to extraction tubes and sterilizing (Fig. S2, transfer), five surface sterilization controls of the fluid used to wash specimens prior to dissection (Fig. S2, wash), and six extraction kit controls from both the DNeasy and powersoil kits (Fig. S2, dneasy and powersoil).

### 16S rRNA Amplicon Data Processing

We denoised and merged reverse and forward reads in QIIME 2 (51) using the DADA2 pipeline (52) separately for each dataset, with the exception of the B3 dataset in which we only considered forward reads due to high error rates in the reverse reads. We aligned the denoised sequences in MAFFT (53), filtered the alignments, and constructed a midpoint-rooted phylogeny using the “align-to-tree-mafft-fasttree” pipeline in QIIME 2. We classified amplicon sequence variants (ASVs) with the QIIME 2 “classify-sklearn” plug-in after training a classifier with the “fit-classifier-naive-bayes” plug-in on sequences of the V4 region of 16S rRNA extracted from SILVA 99% OTUs (Release 132) database. The resulting feature tables, phylogenies, classifications, and sample metadata were primarily processed using the R package *phyloseq* (54). We removed ASVs classified to mitochondria, chloroplast, cicada bacterial endosymbiont (*Hodgkinia* and *Sulcia*), eukaryote, and archaea. To minimize noise introduced by sequencing errors, unclassified taxa, and low sequencing coverage, we removed ASVs that could not be classified at the Kingdom, Phylum, or Class taxonomic levels and ASVs with a total abundance across all samples within a dataset of less than three. We included only samples with a total abundance greater than 100. The remaining ASVs per dataset were further collapsed using the “tip_glom” function in *phyloseq* to cluster ASVs based on cophenetic distances with a tree height of 0.03. We then used the R package *decontam* (55) to identify putative contaminants. However, we were unsuccessful in identifying contaminants with our post-PCR DNA concentration data using the frequency-based filters under reasonable thresholds due to large inter-sample variability in the presence and abundance of different ASVs. Instead, we relied on control samples in the B2 and B3 datasets. We used the prevalence-based filter with a threshold of 0.3 to remove ASVs that were enriched in the controls and then we subsequently removed all ASVs in all datasets that were classified to these putative contaminant genera (Fig. S2).

### Quantitative PCR (qPCR) of 16S rRNA amplicons

To estimate how initial copy numbers of target molecules may differ across our samples, we submitted a subset of samples for qPCR using universal V4 region primers and a CFX Opus 384 Real-Time PCR System. We sought to assess whether quantification of samples after PCR but before pooling would be predictive of the number of target molecules initially present. Thus we analyzed DNA extracts of 24 samples including those of six negative controls and other samples with a range of expected amplicon copy numbers based upon initial quantification as well as sequencing results (Fig. S3 & Table S4). Samples were quantified in triplicate and average initial amplicon copy number estimated based upon correlation to curve calculated with standard.

### Cicada Mitochondrial Genome Phylogeny

We assembled host mitochondrial genomes using off-target capture data from an anchored hybrid enrichment dataset of worldwide cicada lineages (56, 57). We first deduplicated merged and unmerged reads using the function clumpify in BBMap (58), trimmed adapters and reads with Q < 20 using Trimmomatic (59), and then assembled reads using both merged and unmerged data in SPAdes v. 3.12.0 (60, 61). We extracted mitochondrial contigs from the resulting assembly using tblastn with a published partial *K. muta* reference mitochondrial genome – Genbank MG737737 (62) – used BWA v. 0.7.5a (63) in a second processing step, and then reassembled the resulting reads with SPAdes 3.12.0. The final sets of mitochondrial contigs for each sample were aligned to various published and nearly complete mitochondrial genomes (62) using the MAFFT v. 7 E-INS-i algorithm (64), combined into single mitochondrial genome sequences, and manually edited to exclude misassembled regions in Geneious v. 10.1.3 (65). We used these mitochondrial sequences as sample-specific baits in MITObim v. 1.9.1 (66) to assemble improved, higher-quality mitochondrial genomes that were aligned with the MAFFT v. 7 E-INS-i algorithm and manually edited in Geneious v. 10.1.3 for a final mitochondrial genome alignment. We designed a partitioning scheme that included combined 1st and 2nd codon positions and the 3rd codon position for each protein-coding gene, partitions for each of the 12S rRNA and 16S rRNA loci, and a single partition for all tRNAs. We ran a maximum likelihood analysis with this partitioning scheme in RAxML v.8 (67) on the CIPRES web server (68) to produce a final phylogeny.

### *Ophiocordyceps* 16S rRNA Amplicons

We annotated *Ophiocordyceps* mitochondrial ASVs manually using *blastn* hits of mitochondrial ASVs against the NCBI nucleotide database and aligned them to the V4 region of 16S rRNA from previously published metagenomes (23). We assembled these metagenomes from raw data downloaded from NCBI’s SRA database using the same assembly pipeline described above (“Cicada Mitochondrial Phylogeny”) and extracted contigs with *blastn* hits to a reference *Ophiocordyceps* 16S rRNA sequence from an annotated mitochondrial genome (*Ophiocordyceps sinensis*, Accession: NC_034659.1). We aligned these contigs to our *Ophiocordyceps* ASV sequences and trimmed the alignment to the V4 region of 16S rRNA. This alignment was used to construct a PCA of all *Ophiocordyceps* ASVs in this study with the homologous sequences from the re-assembled *Ophiocordyceps* cicada symbionts and *Ophiocordyceps sinensis*.

## Results

### Patterns of bacterial communities of New Zealand cicadas suggest a low abundance and highly variable microbiota not present in eggs

Prior to filtering each dataset, some samples contained low total ASV abundances (Fig. 1A, top), suggesting that cicada gut tissues sometimes contain low amounts of bacterial cells or that the dissection procedure failed to isolate many microbial cells. Our qPCR data of 16S rRNA amplicons from a subset of our samples corroborate these patterns (some of which were quantified as possessing fewer than 100 initial copies of the 16S ribosomal rRNA) and show that the absolute quantity of 16S rRNA amplicons inferred from qPCR is positively correlated with the DNA concentration data we collected post-PCR for a subset of samples (Fig. S3, R^2 0.5375). Our results also showed that controls were consistently quantified as having very few initial target molecule copies (fewer than 0.5 in all cases). Herein, we use our DNA concentration data to inform our filtering procedure.

**Fig. 1:**
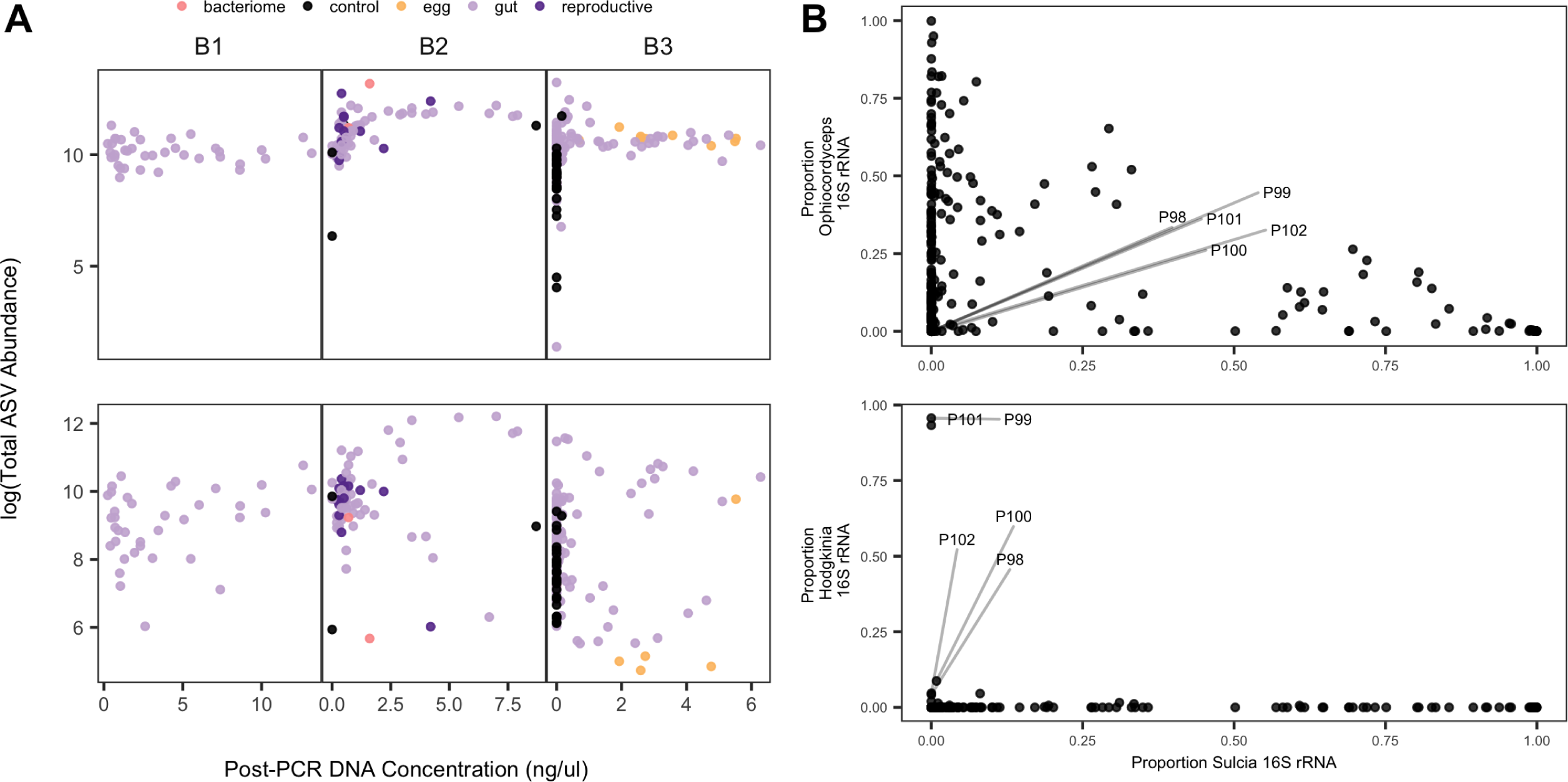
(A) The relationship between DNA concentrations of the amplicon libraries and logged total abundance of sequenced ASVs per sample. The top three panels represent unfiltered data corresponding to each dataset and the bottom three panels represent filtered bacteria-only data corresponding to each dataset. Colors correspond to tissue type. (B) Relationship between the relative abundance of *Sulcia* and either *Hodgkinia* (top) or *Ophiocordyceps* (bottom) across all samples. Labeled points are *Magicicada septendecim* samples.

All datasets contained high relative abundances of *Sulcia*, *Hodgkinia*, or *Ophiocordyceps* ASVs and comparatively few ASV’s of other microbial symbionts (Fig. 1A and Fig. S4). Notably, egg samples in the B3 dataset were nearly entirely composed of *Sulcia* (but contained no *Hodgkinia*) with negligible abundances of other bacteria (Fig. 1A, bottom and Fig. S4). The proportion of *Ophiocordyceps* ASVs negatively scales with the proportion of *Sulcia* ASVs as would other co-obligate symbionts in which both symbionts are present within an individual cicada but one is selectively amplified compared to the other depending on starting quantities (Fig. 1B). This, and the absence of *Hodgkinia*, lends evidence to a scenario in which *Ophiocordyceps* is a widespread resident across our samples.

While *Magicicada* samples contained high relative abundances of *Hodgkinia*, we did not find high abundances of this endosymbiont in any of the New Zealand cicadas, the New Caledonian *Caledopsalta* cicadas (phylogenetic outgroup of the New Zealand cicadas), either of the *Neotibicen* species, or *Platypedia.* We assume small but detectable levels of *Hodgkinia* in non-*Magicicada* samples are due to low levels of contamination from *Magicicada* samples that were concurrently processed (Fig. S5). When *Hodgkinia* does occur as a symbiont in our dataset (i.e., in *Magicicada* samples) it is in high abundance (11, 18, 27, 28). We also found that all taxa, except *M. septendecim* and the *Caledopsalta* specimens, included at least one specimen with high relative abundance *Ophiocordyceps* ASVs (Fig. 2A). Although we did not detect either *Hodgkinia* or *Ophiocordyceps* in the *Caledopsalta* specimens, these samples were relatively old and greater sample sizes are needed to confirm the absence of these two known cicada symbionts in New Caledonian cicadas. The lack of *Hodgkinia* may also be caused by poor primer match due to *Hodgkinia*’s rapid evolution, however this is not compatible with the widespread pattern of *Hodgkinia* absence and *Ophiocordyceps* presence we report in this study on the basis of 16S rRNA (Fig. 2A).

**Fig. 2:**
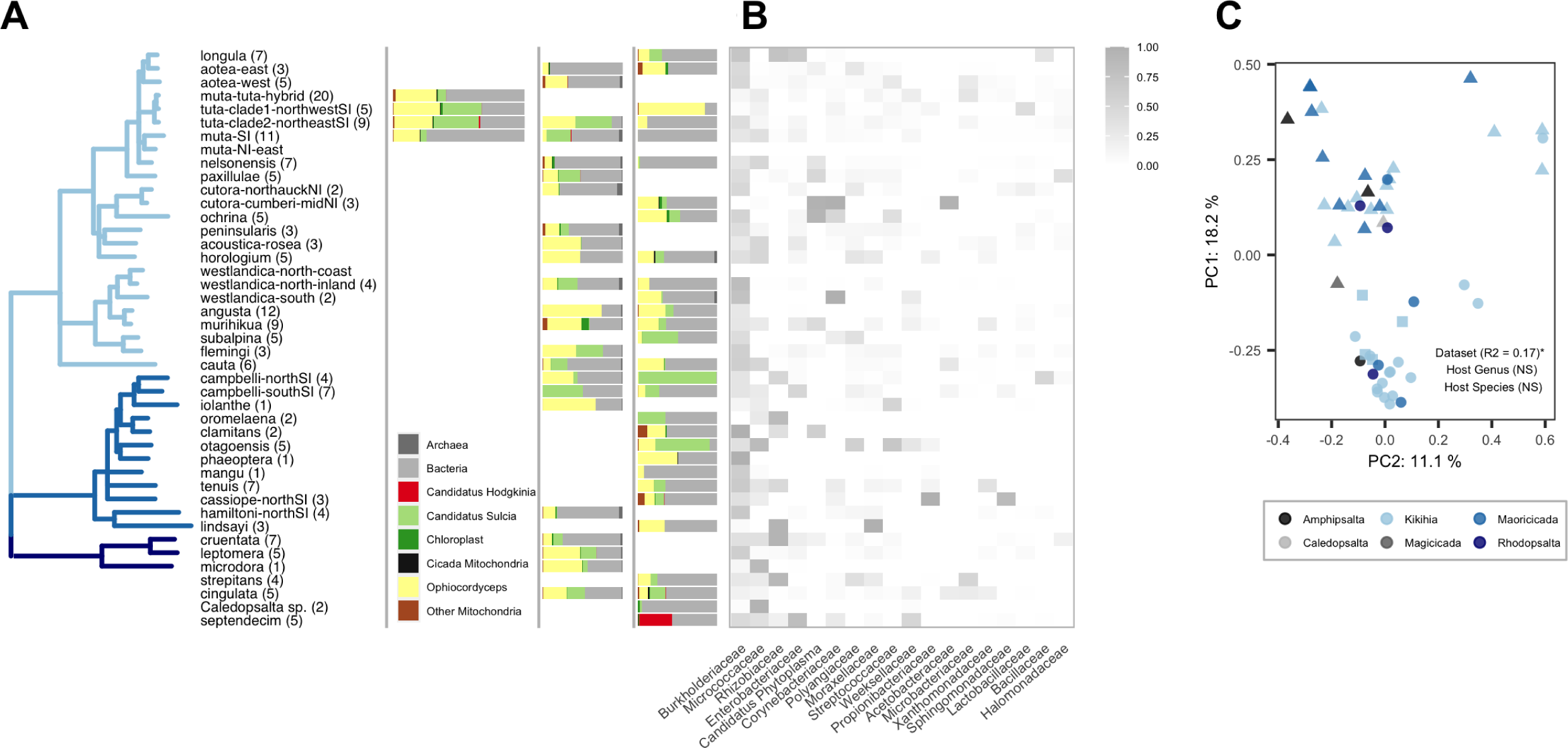
**Distribution of major taxonomic groups using 16S V4 rRNA classification of amplicon sequence variants (ASVs) across all datasets**. **(A)** Maximum likelihood phylogeny using nearly complete mitochondrial genomes of the sampled host species. Branches are collapsed at a bootstrap threshold of 90; bootstrap supports for all nodes are shown in Fig. S11 (see Fig. S11 at https://doi.org/10.6084/m9.figshare.16416720.v1). Tip labels contain the number of individuals sampled in parentheses and match specific epithets in Table S1. Branch colors represent different genera as in A. Species not used to construct the phylogeny are added below the tree (*Amphipsalta strepitans*, *Amphipsalta cingulata*, *Caledopsalta* sp., *Magicicada septendecim*). (Right) The relative abundance of major taxonomic groups averaged across individuals within each host species for dataset B1, dataset B2, and dataset B3. **(B)** Heatmap of relative abundances of major bacterial families averaged across individuals and datasets per species. **(C)** Ordination using weighted UniFrac distances of filtered bacteria-only samples using relative abundances of major bacterial families averaged across individuals within species. Colors correspond to host host genera. Shapes correspond to datasets (square = B1, circle = B2, triangle = B3). The axis labels report the percent variation explained by the corresponding principle components. R-squared values of a permanova analysis using dataset, host genus, and host species as predictors are reported when they were significant at a 5% significance level.

After filtering samples (see “16S rRNA Amplicon Data Processing” in Methods), we identified 18 major bacterial families with high relative abundance and prevalence across samples and datasets, with Burkholderiaceae, Micrococcaceae, Rhizobiaceae (non-*Hodgkinia*), and Enterobacteriaceae among the top four groups in terms of relative abundance (Fig. 2B, S6, see Fig. S7 at https://doi.org/10.6084/m9.figshare.16432914.v1). We could not detect a qualitative structuring of these major bacterial families among clades defined by the host mitochondrial genome phylogeny (Fig. 2B, S6). We used principal coordinates analysis (PCoA) ordinations of Bray distances derived from relative abundances of ASVs belonging to these major bacterial groups and averaged across samples within species (Fig. 2C) to show that variation in bacterial communities surveyed in this study are primarily explained by differences in datasets (permanova, R^2^ = 0.17, p < 0.01) rather than differences in genera or species sampled (permanova, p = NS). In addition, we could not find significant contributions of host genera or species in structuring communities within datasets using either weighted and unweighted UniFrac distances of unmerged samples (see Fig. S8 at https://doi.org/10.6084/m9.figshare.16432929.v1). Although specimens used in this study were collected over many years, we do not find a qualitative effect of collection date in structuring communities (see Fig. S9 at https://doi.org/10.6084/m9.figshare.16432938.v1) and the effect of collection year was insignificant in explaining microbial community differences when considering each dataset separately (permanova, p = NS). After filtering, we were left with one nymphal sample with a similar microbial composition as adult samples (see Fig. S10 at https://doi.org/10.6084/m9.figshare.16432947.v1), but we are unable to make conclusions about the nymphal microbiome with these data alone.

### Mitochondrial genome phylogeny improves resolution of relationships among New Zealand cicadas

Our host phylogeny is based on a whole mitogenome data, nearly seven-fold increase in alignment length from previous phylogenies (69, 70), which were based on approximately 2000 bp of sequence data. Our phylogeny (Fig. 2A, S11) recovers *Rhodopsalta microdora* as sister to *R. leptomera* + *R. cruentata* with 100% bootstrap support, confirming results with fewer genes (71), and groups *Rhodopsalta* with *Maoricicada* with 100% bootstrap support. The *Maoricicada* taxon sampling is reduced compared to the previous *Maoricicada* mitochondrial tree, but despite the missing taxa the mitogenome phylogeny is congruent with the previous tree (69). For the *Kikihia* taxa, the mitogenome phylogeny recovers the same major well-supported clades previously identified (70).

### Elevational differences explain bacterial community variation better than host phylogeny

We used various statistical tests to assess the effect of host phylogeny on bacterial community variation and found no evidence of host phylogenetic structure. Mantel tests between unweighted and weighted UniFrac distances among gut microbial communities and cophenetic distances among host mitochondrial genomes were insignificant based on permutation tests for both B2 and B3 datasets (p >> 0.5). Tests of phylogenetic signal (Pagel’s lambda and K statistic) using the first axis positions of either unweighted or weighted UniFrac PCoA ordinations as phylogenetically distributed traits were insignificant for each dataset as well (p >> 0.5). In addition, ordinations of both weighted and unweighted UniFrac distances did not produce clustering that corresponded to host taxonomy at either the genus or species levels (Fig. 2C and see Fig. S8 at https://doi.org/10.6084/m9.figshare.16432929.v1).

We were able to collect specimens from a wide range of elevations throughout New Zealand (Fig. 3A), allowing us to isolate the effects of phylogenetic and elevational relationships among specimens. We did not find a positive correlation between phylogenetic distance and pairwise bacterial community dissimilarity using either weighted or unweighted UniFrac distances in either the B2 or B3 datasets (Fig. 3B, right), in accordance with the lack of host phylogenetic structure among the most abundant and prevalent bacterial families (Fig. 2B).

**Fig. 3:**
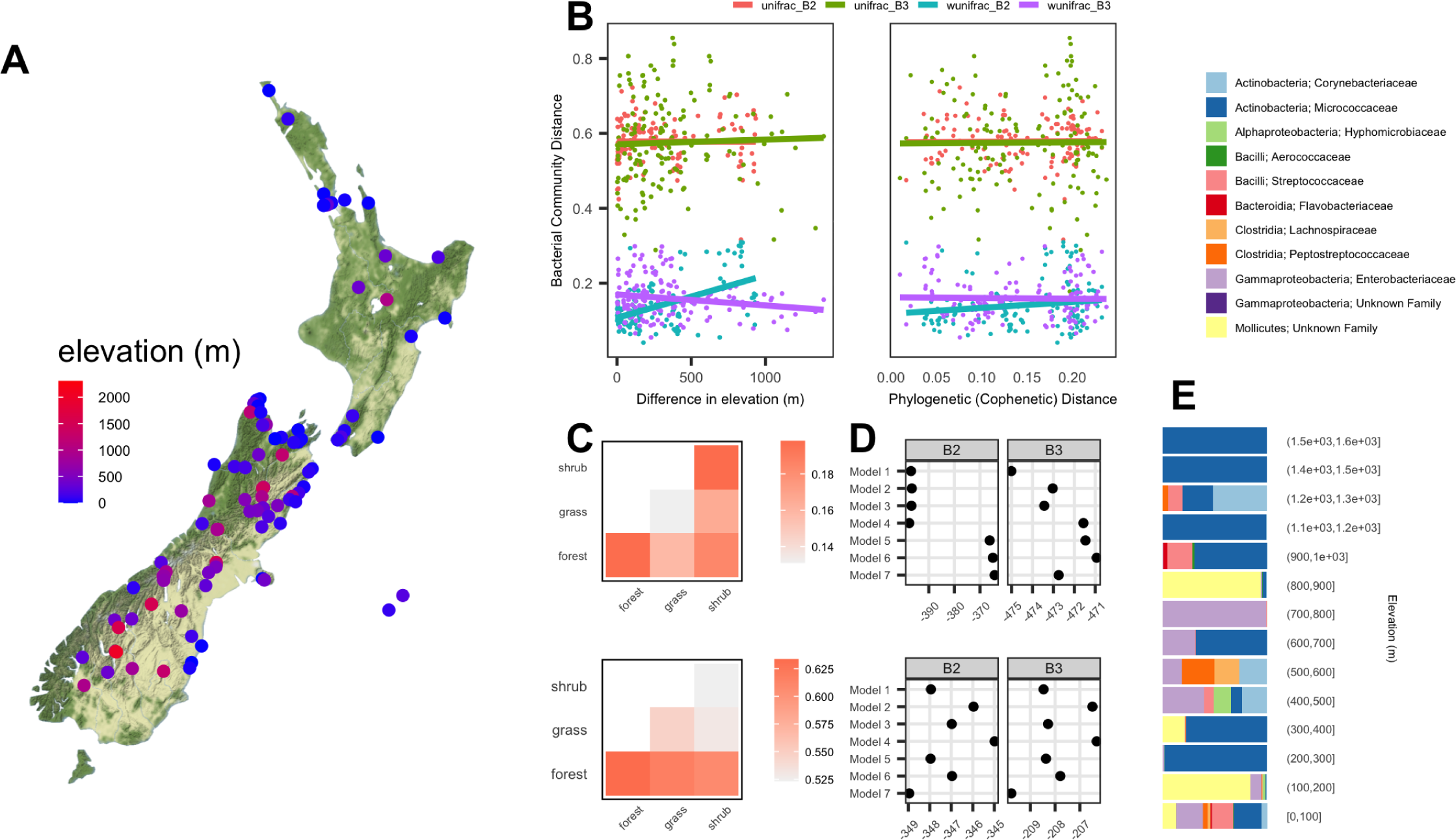
**Correlates of bacterial community differences in New Zealand cicadas using filtered 16s V4 rRNA ASVs from the B2 and B3 datasets**. **(A)** Sampled sites with labels representing state and locality codes found in Table S1. **(B)** Pairwise bacterial community differences using both unweighted (unifrac) and weighted (wunifrac) UniFrac distances with increasing differences in elevation (left) and increasing cophenetic distances calculated using the host mitochondrial phylogeny (right) per dataset. Pairwise comparisons between samples representing different host genera were excluded. **(C)** Average pairwise bacterial community differences among *Kikihia* samples in the B2 dataset using weighted (top) and unweighted (bottom) UniFrac distances for comparisons between samples of the same and different habitats. **(D)** AIC values across beta regression models using weighted (top) and unweighted (bottom) UniFrac distances as response variables. Model formulas, including explanatory variables, associated with each model are provided in Table S2. **(E)** Relative abundance of differentially abundant family-level ASVs across samples binned into increasing elevation categories. Family-level bacterial ASV classifications (above) were found to be significantly differentially abundant between high and low elevation samples based on analysis with DESeq. Note that the group “Mollicutes;Unknown Family” (yellow) is represented entirely by ASVs belonging to the plant pathogen Candidatus *Phytoplasma*.

However, we found a strong positive correlation between differences in elevation and weighted UniFrac pairwise distances in the B2 dataset, suggesting elevational differences may play an important role in structuring New Zealand cicada microbial communities (Fig. 3B, left). In addition to elevation, we examined the effect of habitat type (forest, grassland, or shrub habitats) specifically in *Kikihia* species because these species occupy a broad range of habitat types. Our collections and observations of these species allow us to group them into discrete habitat types (see Fig. S12 at https://doi.org/10.6084/m9.figshare.16416777.v1), however e did not find significant differences between samples of the same habitat type and samples from different habitat types (Fig. 3C)

In addition, we compared multiple Beta regression models in which cophenetic distances between host lineages were used in combination with other covariates or excluded. Both the best and worst fitting models (AIC) included the effect of cophenetic distances, suggesting that host phylogeny had negligible explanatory power compared to the ecological covariates that were also included in these models (Fig. 3D and Table S2). We do not find similarly strong patterns in the B3 dataset, but this dataset was enriched for low-abundance bacteria resulting from nested PCR amplification (see “Amplicon sequencing of 16S V4 rRNA” in Methods) which likely erased signals of a positive relationship between elevation and weighted bacterial community distances given methodological effects on relative abundances. However, we find that in both the B2 and B3 datasets, the best fitting Beta regression models of weighted UniFrac distances always included elevation as an explanatory variable (Fig. 3D, top) and that elevation was significant in all models using the B2 dataset despite having small effect sizes (Table S2). These results were corroborated by multivariate analysis (Table S3).

### Microbial communities in hybrids between *Kikihia muta* and *K.* “tuta” resemble those of their parental species

Specimens with evidence of introgression between *K*. *muta* and *K.* “tuta”, which we refer to as hybrids, in the B1 dataset did not show qualitative differences in the relative abundance of bacterial ASVs compared to parental species (Fig. 4A). Ordination of gut microbial communities using unweighted and weighted UniFrac distances showed that hybrids cluster with parental species (Fig. 4B) with significant effects of processing date (i.e., time at which samples were purified for DNA) and sample depth (i.e., total abundance of all ASVs). The processing date explained 9% and 11% of variation in unweighted and weighted UniFrac distances (permanova, p << 0.05), respectively. However, hybrid status did not significantly explain variation in weighted UniFrac distances (permanova, p >> 0.05) and explained less variation than processing date in unweighted UniFrac distances (permanova, R2 = 7%, p << 0.05).

**Fig. 4:**
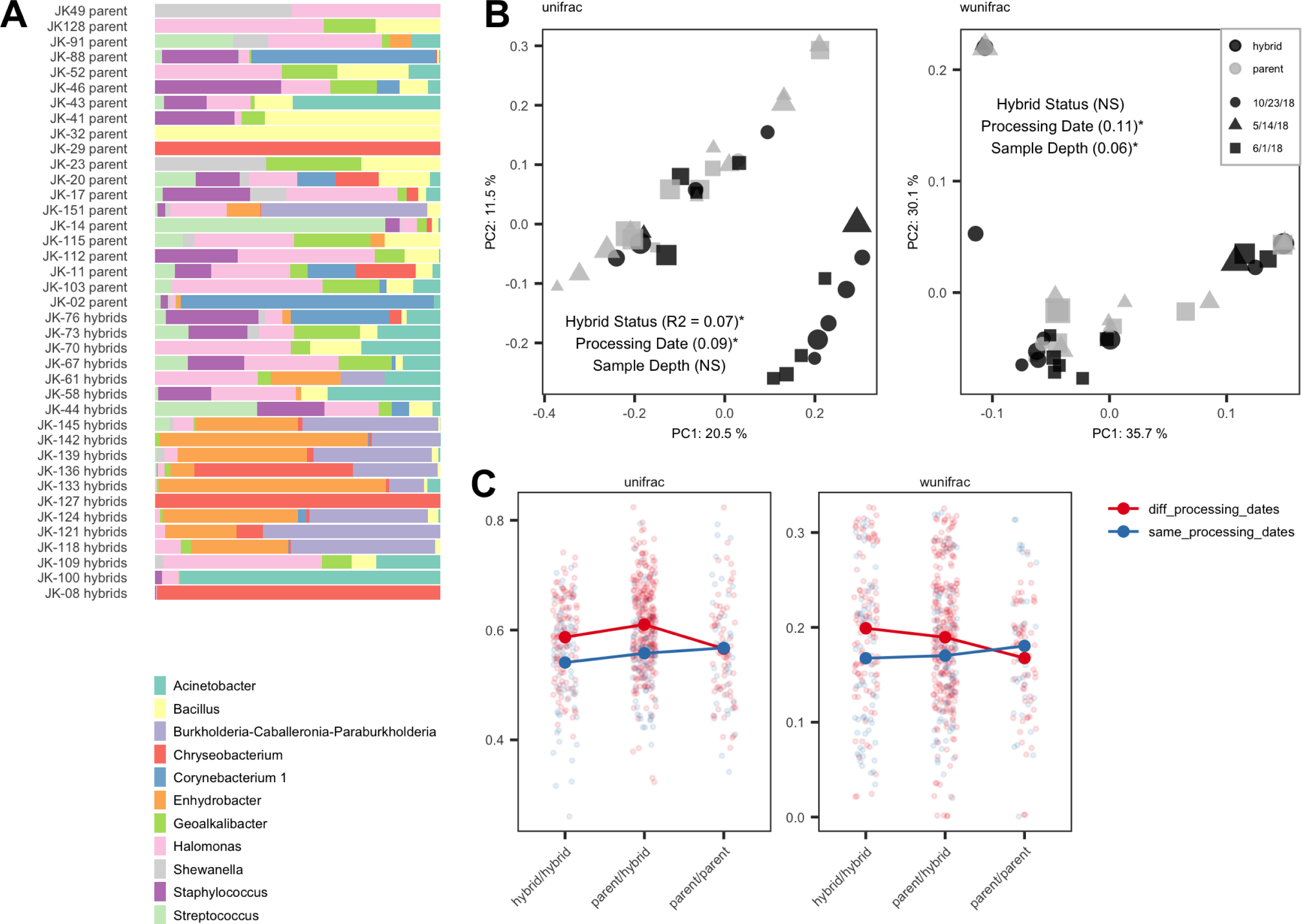
Bacterial community differences among hybrid and parental *Kikihia muta* and *Kikihia tuta* using filtered 16S V4 rRNA ASVs from the B1 dataset. (A) PCoA ordination of samples from dataset B1 using unweighted UniFrac (left) distances and weighted UniFrac distances (right). Grey samples represent hybrid cicadas and black samples represent parental samples of either *Kikihia muta* or *Kikihia tuta*. Shapes represent different times of dissection and sample processing and their size represents relative sample depth (i.e., total ASV abundance). R-squared values from permanova analyses of the effect of hybrid status, processing date, and sample depth on each respective distance measure are reported in parentheses when explanatory variables were significant at a 0.05 significance level. **(B)** Unweighted (left) and weighted (right) pairwise bacterial community distances for hybrid/hybrid, hybrid/parent, and parent/parent sample comparisons. Colors represent comparisons in which samples were processed either within the same processing date or within different processing dates. Lines connect mean values for each category of comparisons. **(C)** Relative abundance of the most abundant Genus-level bacterial ASVs across hybrids and parental samples.

Pairwise unweighted and weighted UniFrac distances were elevated in comparisons of samples that included a hybrid specimen (Fig. 4C, hybrid/hybrid and hybrid/parent pairwise comparisons). Although this pattern may suggest that hybrids have a divergent and more variable microbiome compared to parents, these results are confounded by the effects of processing date. Considering only comparisons of samples processed on the same day (Fig. 4C, same_processing_dates), comparisons between hybrid and parental samples have indistinguishable differences in their bacterial communities. We found that the mean and variance of bacterial community distances are indistinguishable between comparisons that included a hybrid specimen and comparisons that included only parental specimens (Kolmogorov-Smirnov Test, p >> 0.05; F test, p >> 0.05).

### New Zealand cicadas harbor *Ophiocordyceps* instead of *Hodgkinia* but retain *Sulcia*

We annotated all ASVs that were classified to mitochondria as *Ophiocordyceps* fungal symbionts if they matched known *Ophiocordyceps* in BLAST searches against GenBank. We identified three clusters based on a PCA of aligned 16S V4 rRNA sequences (Fig. 5A). We assume that these *Ophiocordyceps* ASVs represent fungal symbionts in the cicadas we sampled because (1) the absence of *Hodgkinia* suggests replacement by an alternative symbiont as shown previously (23) and (2) *Ophiocordyceps* ASVs are identical to previously verified (23) *Ophiocordyceps* symbionts rather than *Ophiocordyceps* pathogens, one species of which we have included in our analysis (Fig. 5A, *Ophiocordyceps sinensis*). Based on the distribution of *Ophiocordyceps* ASV clusters across the host phylogeny (Fig. 5B), we find a qualitative association between these *Ophiocordyceps* ASV clusters and three host mitochondrial clades that correspond to *Kikihia* (Fig. 5B, orange *Ophiocordyceps* ASV cluster), *Maoricicada* (Fig. 5B, yellow *Ophiocordyceps* ASV cluster), and *Rhodopsalta* (Fig. 5B, grey *Ophiocordyceps* ASV cluster), as well as specimens that represent the New Zealand cicada genus *Amphipsalta* but were not included in the mitochondrial tree (Fig. 5B, grey and orange *Ophiocordyceps* ASV clusters). In addition to a qualitative association between *Ophiocordyceps* cluster identity and host phylogenetic group, we find an association with average elevation among specimens of each mitochondrial lineage (Fig. 5B, elevation). This is primarily driven by *Maoricicada* samples of certain species which are found at higher elevations than other New Zealand cicada species.

**Fig. 5:**
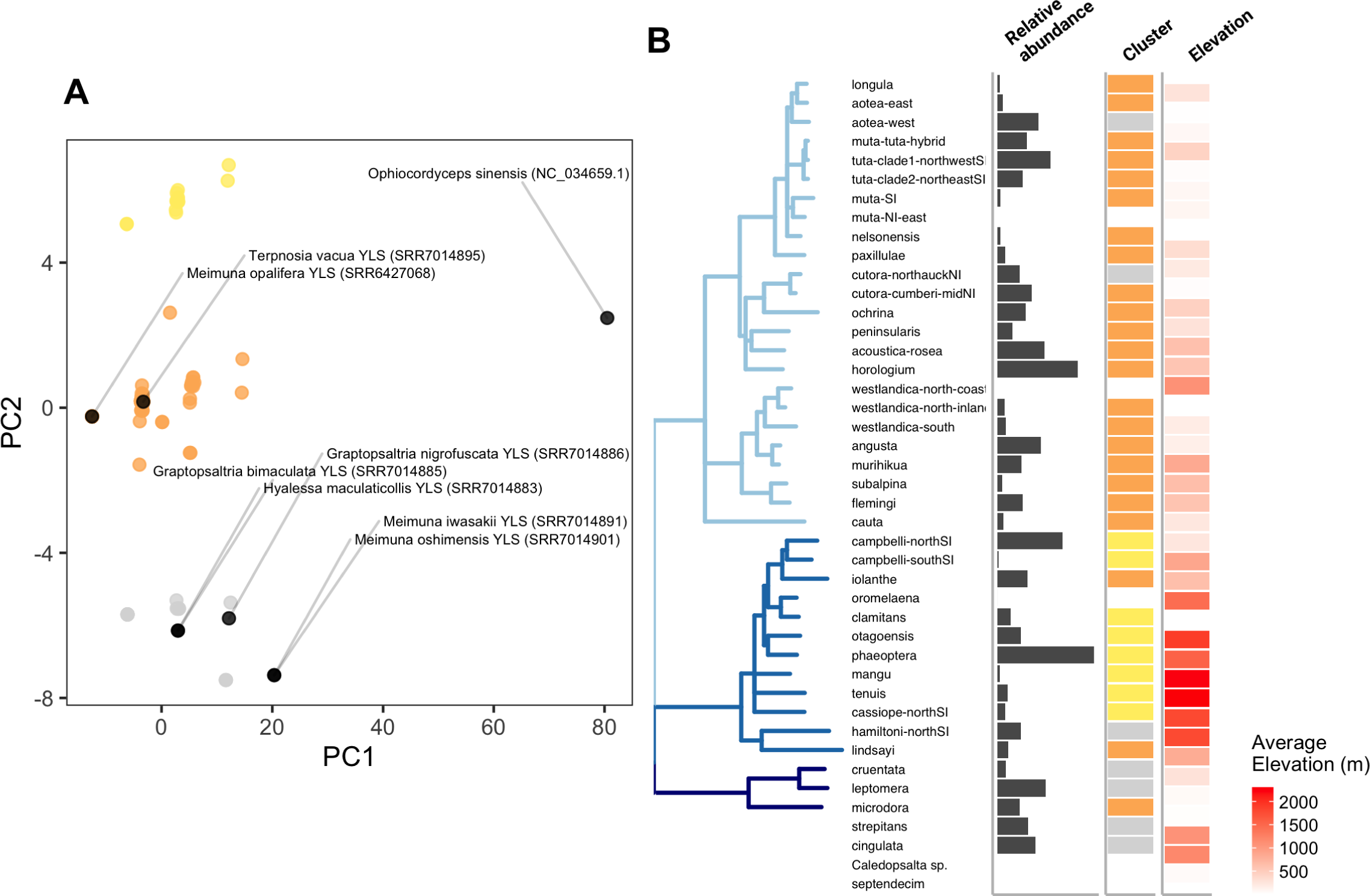
Distribution of *Ophiocordyceps* mitochondrial 16s V4 amplicons and comparison with published *Ophiocordyceps* cicada symbionts across all datasets. (A) PCA of single nucleotide polymorphisms from an alignment of 16S V4 ASV sequences derived in this study with the corresponding genomic sequence extracted from re-assemblies of published cicada metagenomes with verified presence of *Ophiocordyceps* fungal symbionts (Matsuura et al. 2018). The alignment also includes 16S V4 rRNA sequence of an already assembled pathogenic *Ophiocordyceps sinensis*, which is not known to be a beneficial symbiont in cicadas. **(B)** Relative abundance of ASVs classified to *Ophiocordyceps* among all ASVs (left), cluster identity of the most abundant *Ophiocordyceps* ASV based on PCA (middle), and average elevation across all datasets. Host phylogeny is replicated from Figure 2A.

## Discussion

### Lack of stable microbial communities in New Zealand cicadas

Studies of plant sucking bugs with specialized bacteriome-dwelling endosymbionts have with few exceptions (27) not focused on the microbiota outside the bacteriome. Our dataset suggests that adult cicadas have low abundance transient microbial communities in their guts as shown in some other insects (42, 43). Furthermore, a large proportion of the total abundance of 16S ASVs were classified to endosymbiotic bacteria (*Hodgkinia* and *Sulcia*), fungal symbionts (*Ophiocordyceps*), and/or plant pathogens (Candidatus *Phytoplasma*). Cicada endosymbiotic bacteria are housed in specialized organs, have co-evolved with their cicada hosts for millions of years, and have lost many of the key genes required for independent survival and reproduction (16, 19). Fungal symbionts have been shown to be associated with bacteriomes and/or present in the fat bodies (23). Plant pathogens are also known to circulate in the hemolymph and eventually move to the mouth for transmission to plant hosts (72). We expect that the presence of these microbes in gut tissue samples resulted from contamination of bacteriome, fat body or reproductive tissue and hemolymph during dissection and that their high relative abundances can be partly explained by a lack of abundant bacterial cells in the gut lumen (27). We reason that if the bacterial communities of the cicada gut are genuinely low, that even a small amount of contaminating bacterial DNA from other tissues will show up in our gut amplifications.

In some insects – i.e., caterpillars, (43) – the structural simplicity of the gut niche in combination with the rapid pace at which ingested material moves it is suggested to contribute to the lack of resident microbes (i.e., bacteria that are able to replicate and establish at a higher rate than they are lost due to death or excretion). However, the cicada gut niche is structurally more complex and larger in relative volume and especially surface area than that of most other insects, and includes many separate functional compartments (26, 73) that may be amenable to hosting resident microbes. However, the paucity of abundant and consistent microbial residents in this study may be explained by similar factors as in caterpillars, including the physiological pace of food processing driven by plant transpiration (74) and osmotic pressure gradients via the cicada filter chamber (75). These facets of cicada physiology are likely not amenable to microbial colonization as the quick flow of nutritionally poor xylem fluid through the gut may create a harsh environment in which resident bacteria might not be able to establish and proliferate. In addition to being rapidly transported through the cicada alimentary canal, xylem fluid is primarily composed of water with major solutes that include potassium, sodium, calcium, magnesium, chloride, and phosphate (75). This is analogous to the rapid movement of food in the highly alkaline gut environments of caterpillars, which have been reported to lack appreciable evidence of a resident and functional gut microbiota (43). We assume that relatively few microbes other than those adapted to living in xylem vessels (i.e., plant pathogens) can live in this cicada gut environment. Second, the presence of obligate bacterial and fungal symbionts outside of the gut circumvents any obvious need for the functional dependence of adult cicadas on microbe-derived essential nutrients in the gut, reducing the likelihood of selection for life-history or physiological traits that maintain either vertical inheritance or faithful horizontal acquisition of gut microbiota.

While the patterns shown in this study suggest the lack of a consistent resident gut microbiota, low-abundance microbial communities are difficult to detect with the methods we used considering minimal levels of contamination as well as our limited sampling design. In addition, we were unable to detect differences in total bacterial abundance that correlated with host phylogeny or other abiotic correlates we considered. Despite the lack of a consistent gut microbiota across host taxa, it is uncertain what role gut microbial colonization may have in supplementing an already complete cicada diet or providing other transient functions (e.g., immunity to pathogens) as cicadas navigate fluctuating environments. These questions require ecological experiments and tests of fitness. Given the long life of most cicada nymphs (16) compared to mammals, within which gut microbiota are structured by host phylogeny, and the much shorter span of the adult stage (76, 77), it is unlikely that transient microbes are able to establish within adult cicada guts. A greater sampling of nymphal gut microbial diversity and abundance is needed to understand the contribution of environmentally acquired microbiota in the much longer-lived nymphal stage, as well as the microbes that are transmitted to the adult stage from the nymphal stage. While the lack of microbial diversity in eggs dissected from the abdomens of female cicadas in this study suggests a lack of vertical transmission of non-endosymbiotic bacteria, both the overwhelming relative abundance of cicada endosymbiont cells and the lack of other bacterial cells in these samples prevents sufficient detection of low-abundance microbes.

### New Zealand cicada mitochondrial phylogeny compared to previous studies with much less data

A bonus feature of our study is a new phylogeny of the host NZ cicada species using nearly complete mitochondrial genomic data. Support among the NZ cicada species is improved by the inclusion of most mitochondrial genes (see Fig. S11 at https://doi.org/10.6084/m9.figshare.16416720.v1). Relationships within the *Maorocicada* are better supported but otherwise compatible with that of previous studies based on 2,274 bp of mtDNA with slightly larger taxon sampling (69, 78). Relationships involving *M. campbelli* (Fig. 2A, campbelli-northSI and campbelli-southSI) and *M. iolanthe* (Fig. 2A, iolanthe) remain inconsistent with species trees generated from nuclear loci that group these taxa with the other lower elevation species, *M. hamiltoni* (Fig. 2A, hamiltoni-northSI) and *M. lindsayi* (Fig. 2A, lindsayi), in agreement with similarities in male genitalia, mating songs, and habitat specialization (69, 78). Low support at the base of the *Maoricicada* alpine clade suggests rapid diversification during a period of intense mountain building (69). Relationships among *Kikihia* match those of previous mitochondrial phylogenies, including a lack of resolution among some clades despite seven times more data (see Fig. S11 at https://doi.org/10.6084/m9.figshare.16416720.v1). This is not surprising given evidence for extensive hybridization (including mtDNA capture) in this group (50, 79).

### Lack of host specificity in New Zealand cicada microbial communities

Our analysis suggested no host phylogenetic structure among bacterial communities in New Zealand cicadas, even when accounting for variation explained by ecological differences, though several other studies have shown a significant effect of host phylogeny in structuring microbial communities in, for example, corbiculate bees, mammals, tropical birds, ants, deer mice, fruit flies, mosquitoes, and wasps (37, 38, 40, 80, 81). Small phylogenetic distances among host taxa and the suggested lack of functional importance of gut microbial communities likely contributed to these results.

In addition, we could not find significant differences in gut bacterial composition between hybrids and their parental species, reinforcing our conclusions that cicadas lack resident gut microbiota that are structured by host phylogeny. Indeed, if host adaptations that maintain assembly of bacterial communities have evolved, we would expect that these mechanisms in hybrids would deteriorate when genomic variants that have undergone divergent ecological selection for gut microbial community assembly (82) are combined into different genetic backgrounds. Improper regulation of gut bacteria in hybrid hosts may be one source of hybrid lethality, however support for this hypothesis is uncertain (83, 84). We found little evidence that host genetics influenced microbial community assembly, as shown in other hybrid crosses (82, 85). Although gut microbial communities in New Zealand cicadas do not seem to be host specific, the composition of these communities may be highly influenced by elevational differences among host individuals. Elevational diversity is a prominent feature of the New Zealand landscape, with dramatically varying ecosystems along elevational gradients.

Understanding the many ecological factors contributing to variation in the microbiome is an important next step for understanding host-associated microbial diversity. While our data suggest an association with elevation, we lack sufficient power to identify specific bacterial taxa enriched at high or low elevations. In addition, future studies with broader within-species sampling are needed to investigate the causes of intraspecific variation in the microbiome.

Although we do not have sufficient intraspecific sampling of elevationally diverse species in this study, some *Kikihia* in particular occupy broad elevational ranges and may be good candidates for additional sampling (see Fig. S12 at https://doi.org/10.6084/m9.figshare.16416777.v1). Nonetheless, elevation is the best predictor of microbial community variation in our analyses.

### Phytoplasma plant pathogens in New Zealand cicadas

We report the first sequence-based identification, to our knowledge, of the plant pathogen *Candidatus* Phytoplasma in cicadas. Phytoplasmas (Phylum Tenericutes: Class Mollicutes) comprise a group of prokaryotic relatives to mycoplasmas and spiroplasmas and are transmitted among plant hosts by insect vectors primarily represented by the hemipteran groups Cicadellidae, Fulgoromorpha, and Psyllidae, but may be transmitted by the heteropteran families Pentatomidae and Tingidae as well (72). Despite interactions with both plant host and insect vector, phytoplasma strains do not appear to have insect-vector specificity and many different vectors may transmit a single phytoplasma strain or vice versa (72). This pattern is recapitulated in other insect-vectored plant pathogens, particularly *Xylella* pathogens which are vectored by the apache cicada *Diceroprocta apache* into California grapevine (86) and by various Brazilian cicadas into coffee (87). Phytoplasmas were found in high abundance across six specimens representing *K. paxillulae, K.* “murihikua”, *K.* “aotea”, *K. muta*, *K.* “nelsonensis”, and *K. ochrina*. We did not detect phytoplasmas in any other New Zealand cicada genus other than *Kikihia*, which tend to occupy lower elevation habitats characterized by grasses and shrubs (Fig. 2B, S6, see Fig. S7 at https://doi.org/10.6084/m9.figshare.16432914.v1). The high abundance of 16S rRNA amplicons that classified to phytoplasmas after our filtering procedure (i.e., Phytoplasmas comprised a large majority of that sample’s total rRNA abundance when present) suggests that New Zealand cicadas are a new potential vector of these plant pathogens.

### Evidence for endosymbiont replacement by *Ophiocordyceps* fungal symbionts in New Zealand cicadas

We provide partial evidence using mitochondrial 16S rRNA amplicons that New Zealand cicadas (and some North American cicadas, including *Platypedia putnami* and two species of *Neotibicen*) lack *Hodgkinia* bacterial endosymbionts and instead harbor high abundances of *Ophiocordyceps* fungus (Fig. 2A). Cicada taxa enriched for *Ophiocordyceps* mitochondrial 16S rRNA in our study lacked *Hodgkinia* 16S rRNA, mirroring the findings of a previous study of Japanese cicadas (23). Given the lack of adequate sequence data in this study, we cannot resolve the relationships among *Ophiocordyceps* symbiont strains, however we were able to identify signals of genetic structure in mitochondrial 16S rRNA that correlate with the host mitochondrial phylogeny (Fig. 5). In particular, we found that different cicada genera sampled in this study likely have unique *Ophiocordyceps* genotypes based on mitochondrial 16S rRNA.

These putative *Ophiocordyceps* symbionts in *Kikihia, Rhodopsalta*, and *Amphipsalta* were different from those in the higher elevation *Maoricicada*. We found that three *Maoricicada* species had *Ophiocordyceps* genotypes that did not match all other *Maoricicada* and instead clustered with *Kikihia*, *Rhodopsalta*, and *Amphipsalta* genotypes (*M. iolanthe, M. lindsayi,* and *M. hamiltoni*). These are unlikely cases of independent re-acquisitions of *Ophiocordyceps* symbonts. A previously published optimal species tree using nuclear and mitochondrial gene markers (69) placed *M. iolanthe, M. lindsayi*, and *M hamiltoni* together with *M. myersi* as a monophyletic group sister to *M. campbelli*. Considering that *M. iolanthe, M. lindsayi,* and *M. hamiltoni* are lowland *Maoricicada* species, *Ophiocordyceps* symbionts in all other, highland, *Maoricicada* species may have experienced accelerated evolution at the time of the rise of high elevation habitat in New Zealand resulting in a unique *Ophiocordyceps* genotype in highland taxa. Although *M. campbelli* ranges from low to mid elevation and shares genitalic characteristics with *M. iolanthe, M. lindsayi,* and *M. hamiltoni*, its *Ophiocordyceps* genotype groups with other, non-lowland, species.

Although we did not find evidence of either *Hodgkinia* or *Ophiocordyceps* in the sister-group New Caledonian cicadas, additional sampling is needed to verify this result. In addition, sampling of Australian species, which are closely related to the clade formed by the New Zealand cicada genera *Amphipsalta* and *Notopsalta,* will also be required to resolve the question of whether *Ophiocordyceps* symbionts were acquired before or after cicadas colonized New Zealand. NZ cicadas are not parasitized by *Ophiocordyceps* but rather a fungus in the genus *Isaria* (88). Other NZ insects, however, are parasitized by *Ophiocordyceps*. Indeed, the number and extent of independent *Ophiocordyceps* domestications in cicadas requires greater sampling of fungal pathogen diversity as well.

## Conclusion

In summary, we find that gut microbial communities in NZ cicadas are not strongly structured by host phylogeny and that hybrid cicadas resemble parental species, suggesting a lack of host specificity. Instead, we find that elevation differences among individuals explain variation in gut microbiota better than phylogenetic relatedness among hosts. Our new and nearly complete mitochondrial genomes improved phylogenetic relationships among NZ cicadas, providing a valuable resource for investigations of the evolutionary history of this group. In addition, we have shown what is likely the loss of the obligate symbiont *Hodgkinia* and the acquisition of *Ophiocordyceps* symbionts in NZ cicadas. This pattern has previously only been shown in 13 genera of cicadas collected in Japan (nine of which possess *Ophiocordyceps* and three of which possess *Hodgkinia*) and three previously-studied genera from the Americas which all possess *Hodgkinia* (23); These represent four lineages from the cicada subfamily Cicadinae, four lineages from the family Cicaettinae and one from the subfamily Tibicinninae, all of which are widely distributed outside of Japan (89-91). Our study (which is the first to find *Ophiocordyceps* replacing *Hodgkinia* in the southern hemisphere) together with these previous insights suggest that replacements of obligate bacterial symbionts in Cicadoidea are more common and widespread than previously thought and that they may proceed via convergent domestications of similar fungal pathogens. A broad survey of cicada endosymbionts worldwide is in progress in our laboratory and others. Together, these findings highlight the instability of insect-microbe associations (92) and show that instability is present at different time scales, from the rapid ecological turnover of gut microbial communities over host generations to transitions in obligate symbiont identity over millions of years of host evolution.

## Acknowledgements

We would like to thank David C. Marshall for collecting specimens and providing valuable advice during the course of this study. We would like to thank the Microbial Analysis, Resources, and Services (MARS) Facility at the University of Connecticut for help on experimental design and amplicon sequencing of microbial communities, the University of Connecticut’s High Performance Computing (HPC) Facility for providing computational resources for data processing and analysis, the New Zealand Department of Conservation for help in obtaining permits to collect cicadas, and the Society for the Study of Evolution for a travel grant to present this work at the Evolution 2019 conference. We would like to thank funding resources from NSF GRFP (D.H). This study was funded by NSF DEB 1655891 (C.S.), DEB-0955849 (C.S.), DEB 0720664 (C.S.), DEB 0529679 (C.S.), DEB 0422386 (C.S.), DEB 0089946 (C.S.), grants from the University of Connecticut, The New Zealand Marsden Fund, The National Geographic Society and the Fulbright Foundation.

## Supplementary Tables

**Table S1:** Sample metadata

**Table S2:** Results of Beta regressions using either unweighted or weighted pairwise UniFrac distances as response variables and different combinations of explanatory variables (elevation, phylogeny, and habitat) across seven independently run models. Values represent effect sizes with standard errors in parentheses. Asterisks are included when effect sizes are significant (p << 0.01).

**Table S3:** Results of PERMANOVA analysis of the effect of various explanatory variables (elevation, species, and habitat) on weighted UniFrac distances among gut microbial communities across datasets.

**Table S4:** General 16S V4 rRNA qPCR data and associated data from 16S V4 rRNA amplicon sequencing.

## Supplementary Figures

**Fig. S1:** Image of gut tissue dissected from nymphal *Magicicada septendecim*. (A) Complete gut. (B) Filter chamber. (C) Midgut. (D-E) Malphigian tubules and hindgut. (F) Rectum. Adult gut anatomy is identical to that of the nymph in these hemimetabolous insects and varies little across the family.

**Fig. S2:** Relative abundance of a subset of putative contaminants identified with Decontam. Sample labels include control type (powersoil, transfer, wash, dissection, dneasy, and pcr), the dataset in which the control was produced (B2 or B3), and the total abundance of all ASVs before dataset filtering.

**Fig. S3:** The relationship between total DNA post-PCR DNA concentration and qPCR absolute abundance of 16S rRNA in a subset of samples spanning different datasets. Data based on amplification with the same 16S V4 rRNA primers used to sequence microbial communities.

**Fig. S4:** Relative abundance of major taxonomic groups across specimens. Specimen IDs include specific epithets and sample IDs corresponding to Table S1.

**Fig. S5:** (Top) PCA ordination of *Hodgkinia* ASV sequences found across all samples. Colors correspond to ASV clusters representing similar ASV sequences. (Bottom) Abundance of *Hodgkinia* ASV clusters relative to all other *Hodgkinia* ASV clusters across samples.

**Fig. S6:** Relative abundance of major bacterial genera across specimens after dataset filtering. Solid black cells correspond to a relative abundance of 100% and white cells indicate absence of any particular bacterial group.

## Literature Cited

1. McFall-Ngai M, Hadfield MG, Bosch TCG, Carey HV, Domazet-Lošo T, Douglas AE, Dubilier N, Eberl G, Fukami T, Gilbert SF, Hentschel U, King N, Kjelleberg S, Knoll AH, Kremer N, Mazmanian SK, Metcalf JL, Nealson K, Pierce NE, Rawls JF, Reid A, Ruby EG, Rumpho M, Sanders JG, Tautz D, Wernegreen JJ. 2013. Animals in a bacterial world, a new imperative for the life sciences. Proceedings of the National Academy of Sciences 110:3229–3236.

2. Archibald JM. 2015. Endosymbiosis and Eukaryotic Cell Evolution. Curr Biol 25:R911–21.

3. Moran NA. 2007. Symbiosis as an adaptive process and source of phenotypic complexity. Proc Natl Acad Sci U S A 104 Suppl 1:8627–8633.

4. Hurst GDD. 2017. Extended genomes: Symbiosis and evolution. Interface Focus.

5. Moran NA, McCutcheon JP, Nakabachi A. 2008. Genomics and evolution of heritable bacterial symbionts. Annu Rev Genet 42:165–190.

6. Kikuchi Y, Hosokawa T, Fukatsu T. 2007. Insect-microbe mutualism without vertical transmission: a stinkbug acquires a beneficial gut symbiont from the environment every generation. Appl Environ Microbiol 73:4308–4316.

7. Kikuchi Y, Hosokawa T, Fukatsu T. 2011. An ancient but promiscuous host-symbiont association between *Burkholderia* gut symbionts and their heteropteran hosts. ISME J 5:446–460.

8. Hu Y, Sanders JG, Łukasik P, D’Amelio CL, Millar JS, Vann DR, Lan Y, Newton JA, Schotanus M, Kronauer DJC, Pierce NE, Moreau CS, Wertz JT, Engel P, Russell JA. 2018. Herbivorous turtle ants obtain essential nutrients from a conserved nitrogen-recycling gut microbiome. Nat Commun 9.

9. Salem H, Bauer E, Kirsch R, Berasategui A, Cripps M, Weiss B, Koga R, Fukumori K, Vogel H, Fukatsu T, Kaltenpoth M. 2017. Drastic Genome Reduction in an Herbivore’s Pectinolytic Symbiont. Cell 1–12.

10. Bennett GM, Moran NA. 2015. Heritable symbiosis: The advantages and perils of an evolutionary rabbit hole. Proceedings of the National Academy of Sciences 112:10169– 10176.

11. Campbell MA, Łukasik P, Meyer MC, Buckner M, Simon C, Veloso C, Michalik A, McCutcheon JP. 2018. Changes in Endosymbiont Complexity Drive Host-Level Compensatory Adaptations in Cicadas. MBio 9:e02104–18.

12. Buchner P. 1965. Symbiosis in animals which suck plant juices. Endosymbiosis of animals with plant microorganisms 210:432.

13. McCutcheon JP, McDonald BR, Moran NA. 2009. Convergent evolution of metabolic roles in bacterial co-symbionts of insects. Proceedings of the National Academy of Sciences 106:15394–15399.

14. Christensen H, Fogel ML. 2011. Feeding ecology and evidence for amino acid synthesis in the periodical cicada (*Magicicada*). J Insect Physiol 57:211–219.

15. McCutcheon JP, McDonald BR, Moran NA. 2009. Origin of an alternative genetic code in the extremely small and GC-rich genome of a bacterial symbiont. PLoS Genet 5:e1000565.

16. Campbell MA, Van Leuven JT, Meister RC, Carey KM, Simon C, McCutcheon JP. 2015. Genome expansion via lineage splitting and genome reduction in the cicada endosymbiont *Hodgkinia*. Proceedings of the National Academy of Sciences 112:10192–10199.

17. Moran NA, Tran P, Gerardo NM. 2005. Symbiosis and insect diversification: an ancient symbiont of sap-feeding insects from the bacterial phylum Bacteroidetes. Appl Environ Microbiol 71:8802–8810.

18. Campbell MA, Łukasik P, Simon C, McCutcheon JP. 2017. Idiosyncratic Genome Degradation in a Bacterial Endosymbiont of Periodical Cicadas. Curr Biol 0.

19. Łukasik P, Nazario K, Van Leuven JT, Campbell MA, Meyer M, Michalik A, Pessacq P, Simon C, Veloso C, McCutcheon JP. 2018. Multiple origins of interdependent endosymbiotic complexes in a genus of cicadas. Proceedings of the National Academy of Sciences 115:E226–E235.

20. Van Leuven JTT, Meister RCC, Simon C, McCutcheon JP. 2014. Sympatric Speciation in a Bacterial Endosymbiont Results in Two Genomes with the Functionality of One. Cell 158:1270–1280.

21. Müller HJ. 1962. Neuere vorstellungen über verbreitung und phylogenie der endosymbiosen der zikaden. Z Morphol Oekol Tiere 190–210.

22. Müller HJ. 1949. Zur systematik und phylogenie der zikaden-endosymbiosen. Biol Zent 343–368.

23. Matsuura Y, Moriyama M, Łukasik P, Vanderpool D, Tanahashi M, Meng X-Y, McCutcheon JP, Fukatsu T. 2018. Recurrent symbiont recruitment from fungal parasites in cicadas. Proceedings of the National Academy of Sciences 115:E5970–E5979.

24. Noda H. 1977. Histological and histochemical observation of intracellular yeastlike symbionts in the fat body of the smaller brown planthopper, *Laodelphax striatellus* (Homoptera: Delphacidae). Appl Entomol Zool.

25. Hongoh Y, Ishikawa H. 2000. Evolutionary studies on uricases of fungal endosymbionts of aphids and planthoppers. J Mol Evol 51:265–277.

26. Zhou W, Nan X, Zheng Z, Wei C, He H, Goodacre S. 2015. Analysis of inter-individual bacterial variation in gut of cicada *Meimuna mongolica* (Hemiptera: Cicadidae). J Insect Sci 15:1–6.

27. Zheng Z, Wang D, He H, Wei C. 2017. Bacterial diversity of bacteriomes and organs of reproductive, digestive and excretory systems in two cicada species (Hemiptera: Cicadidae). PLoS One 12:1–21.

28. Wang D, Huang Z, He H, Wei C. 2018. Comparative analysis of microbial communities associated with bacteriomes, reproductive organs and eggs of the cicada *Subpsaltria yangi*. Arch Microbiol 200:227–235.

29. Dillon RJ, Dillon VM. 2004. The Gut Bacteria of Insects: Nonpathogenic Interactions. Annu Rev Entomol 49:71–92.

30. Ng SH, Stat M, Bunce M, Simmons LW. 2018. The influence of diet and environment on the gut microbial community of field crickets. Ecol Evol 8:4704–4720.

31. Nishida AH, Ochman H. 2018. Rates of gut microbiome divergence in mammals. Mol Ecol 27:1884–1897.

32. Douglas AE, Werren JH. 2016. Holes in the Hologenome: Why Host-Microbe Symbioses Are Not Holobionts. MBio 7:e02099.

33. Grueneberg J, Engelen AH, Costa R, Wichard T. 2016. Macroalgal Morphogenesis Induced by Waterborne Compounds and Bacteria in Coastal Seawater. PLoS One 11:e0146307.

34. Lin JD, Lemay MA, Parfrey LW. 2018. Diverse Bacteria Utilize Alginate Within the Microbiome of the Giant Kelp *Macrocystis pyrifera*. Front Microbiol 9:1914.

35. Coon KL, Vogel KJ, Brown MR, Strand MR. 2014. Mosquitoes rely on their gut microbiota for development. Mol Ecol 23:2727–2739.

36. Coon KL, Brown MR, Strand MR. 2016. Mosquitoes host communities of bacteria that are essential for development but vary greatly between local habitats. Mol Ecol 25:5806–5826.

37. Kwong WK, Medina LA, Koch H, Sing KW, Soh EJY, Ascher JS, Jaffé R, Moran NA. 2017. Dynamic microbiome evolution in social bees. Science Advances 3:1–17.

38. Brooks AW, Kohl KD, Brucker RM, van Opstal EJ, Bordenstein SR. 2016. Phylosymbiosis: Relationships and Functional Effects of Microbial Communities across Host Evolutionary History. PLoS Biol 14:1–29.

39. Kropáčková L, Těšický M, Albrecht T, Kubovčiak J, Čížková D, Tomášek O, Martin J-F, Bobek L, Králová T, Procházka P, Kreisinger J. 2017. Codiversification of gastrointestinal microbiota and phylogeny in passerines is not explained by ecological divergence. Mol Ecol 26:5292–5304.

40. Hird SM, Sánchez C, Carstens BC, Brumfield R. 2015. Comparative gut microbiota of 59 neotropical bird species. Front Microbiol 6.

41. Hu Y, ??ukasik P, Moreau CS, Russell JA. 2014. Correlates of gut community composition across an ant species (*Cephalotes varians*) elucidate causes and consequences of symbiotic variability. Mol Ecol 23:1284–1300.

42. Hammer TJ, Sanders JG, Fierer N. 2019. Not all animals need a microbiome. FEMS Microbiol Lett https://doi.org/10.1093/femsle/fnz117.

43. Hammer TJ, Janzen DH, Hallwachs W, Jaffe SP, Fierer N. 2017. Caterpillars lack a resident gut microbiome. Proceedings of the National Academy of Sciences 114:9641– 9646.

44. Shapira M. 2016. Gut Microbiotas and Host Evolution: Scaling Up Symbiosis. Trends Ecol Evol 31:539–549.

45. Marshall DC, Hill KBR, Moulds M, Vanderpool D, Cooley JR, Mohagan AB, Simon C. 2016. Inflation of Molecular Clock Rates and Dates: Molecular Phylogenetics, Biogeography, and Diversification of a Global Cicada Radiation from Australasia (Hemiptera: Cicadidae: Cicadettini). Syst Biol 65:16–34.

46. Lane DH. 1995. The recognition concept of speciation applied in an analysis of putative hybridization in New Zealand cicadas of the genus Kikihia (Insects: Hemiptera: Tibicinidae). Speciation and the Recognition Concept: Theory and Application The Johns Hopkins Univ Press, Baltimore, MD.

47. Cooley JR, Marshall DC. 2001. Sexual signaling in periodical cicadas, *Magicicada* spp. (Hemiptera: Cicadidae). Behaviour 138:827–855.

48. Fleming CA. 1975. Adaptive radiation in New Zealand cicadas. American Philosophical Society.

49. Dugdale JS, Fleming CA. 1978. New zealand cicadas of the genus *Maoricicada* (Homoptera: Tibicinida). N Z J Zool 5:295–340.

50. Marshall DC, Hill KBR, Cooley JR, Simon C. 2011. Hybridization, mitochondrial DNA phylogeography, and prediction of the early stages of reproductive isolation: lessons from New Zealand cicadas (genus *Kikihia*). Syst Biol 60:482–502.

51. Bolyen E, Rideout JR, Dillon MR, Bokulich NA, Abnet C, Al-Ghalith GA, Alexander H, Alm EJ, Arumugam M, Asnicar F, Bai Y, Bisanz JE, Bittinger K, Brejnrod A, Brislawn CJ, Titus Brown C, Callahan BJ, Caraballo-Rodríguez AM, Chase J, Cope E, Da Silva R, Dorrestein PC, Douglas GM, Durall DM, Duvallet C, Edwardson CF, Ernst M, Estaki M, Fouquier J, Gauglitz JM, Gibson DL, Gonzalez A, Gorlick K, Guo J, Hillmann B, Holmes S, Holste H, Huttenhower C, Huttley G, Janssen S, Jarmusch AK, Jiang L, Kaehler B, Kang KB, Keefe CR, Keim P, Kelley ST, Knights D, Koester I, Kosciolek T, Kreps J, Langille MGI, Lee J, Ley R, Liu Y-X, Loftfield E, Lozupone C, Maher M, Marotz C, Martin BD, McDonald D, McIver LJ, Melnik AV, Metcalf JL, Morgan SC, Morton J, Naimey AT, Navas-Molina JA, Nothias LF, Orchanian SB, Pearson T, Peoples SL, Petras D, Preuss ML, Pruesse E, Rasmussen LB, Rivers A, Michael S Robeson II, Rosenthal P, Segata N, Shaffer M, Shiffer A, Sinha R, Song SJ, Spear JR, Swafford AD, Thompson LR, Torres PJ, Trinh P, Tripathi A, Turnbaugh PJ, Ul-Hasan S, van der Hooft JJJ, Vargas F, Vázquez-Baeza Y, Vogtmann E, von Hippel M, Walters W, Wan Y, Wang M, Warren J, Weber KC, Williamson CHD, Willis AD, Xu ZZ, Zaneveld JR, Zhang Y, Zhu Q, Knight R, Gregory Caporaso J. 2018. QIIME 2: Reproducible, interactive, scalable, and extensible microbiome data science. e27295v2. PeerJ Preprints.

52. Callahan BJ, McMurdie PJ, Rosen MJ, Han AW, Johnson AJA, Holmes SP. 2016. DADA2: High-resolution sample inference from Illumina amplicon data. Nat Methods 13:581–583.

53. Katoh K, Standley DM. 2013. MAFFT multiple sequence alignment software version 7: improvements in performance and usability. Mol Biol Evol 30:772–780.

54. McMurdie PJ, Holmes S. 2013. phyloseq: an R package for reproducible interactive analysis and graphics of microbiome census data. PLoS One 8:e61217.

55. Davis NM, Proctor D, Holmes SP, Relman DA, Callahan BJ. 2018. Simple statistical identification and removal of contaminant sequences in marker-gene and metagenomics data. Microbiome 6.

56. Lemmon AR, Emme SA, Lemmon EM. 2012. Anchored hybrid enrichment for massively high-throughput phylogenomics. Syst Biol 61:727–744.

57. Simon C, Gordon ERL, Moulds MS, Cole JA, Haji D, Lemmon AR, Lemmon EM, Kortyna M, Nazario K, Wade EJ, Meister RC, Goemans G, Chiswell SM, Pessacq P, Veloso C, Mccutcheon JP, Łukasik P. 2019. Off-target capture data, endosymbiont genes and morphology reveal a relict lineage that is sister to all other singing cicadas. Biol J Linn Soc Lond https://doi.org/10.1093/biolinnean/blz120.

58. Bushnell B. 2014. BBMap: A Fast, Accurate, Splice-Aware Aligner. LBNL-7065E. Lawrence Berkeley National Lab. (LBNL), Berkeley, CA (United States).

59. Bolger AM, Lohse M, Usadel B. 2014. Trimmomatic: a flexible trimmer for Illumina sequence data. Bioinformatics 30:2114–2120.

60. Bankevich A, Nurk S, Antipov D, Gurevich AA, Dvorkin M, Kulikov AS, Lesin VM, Nikolenko SI, Pham S, Prjibelski AD, Pyshkin AV, Sirotkin AV, Vyahhi N, Tesler G, Alekseyev MA, Pevzner PA. 2012. SPAdes: a new genome assembly algorithm and its applications to single-cell sequencing. J Comput Biol 19:455–477.

61. Nurk S, Bankevich A, Antipov D, Gurevich A, Korobeynikov A, Lapidus A, Prjibelsky A, Pyshkin A, Sirotkin A, Sirotkin Y, Stepanauskas R, McLean J, Lasken R, Clingenpeel SR, Woyke T, Tesler G, Alekseyev MA, Pevzner PA. 2013. Assembling Genomes and Mini-metagenomes from Highly Chimeric Reads, p. 158–170. In Research in Computational Molecular Biology. Springer Berlin Heidelberg.

62. Łukasik P, Chong RA, Nazario K, Matsuura Y, Bublitz DAC, Campbell MA, Meyer MC, Van Leuven JT, Pessacq P, Veloso C, Simon C, McCutcheon JP. 2019. One Hundred Mitochondrial Genomes of Cicadas. J Hered 110:247–256.

63. Li H, Durbin R. 2009. Fast and accurate short read alignment with Burrows-Wheeler transform. Bioinformatics 25:1754–1760.

64. Katoh K, Rozewicki J, Yamada KD. 2017. MAFFT online service: multiple sequence alignment, interactive sequence choice and visualization. Brief Bioinform https://doi.org/10.1093/bib/bbx108.

65. Kearse M, Moir R, Wilson A, Stones-Havas S, Cheung M, Sturrock S, Buxton S, Cooper A, Markowitz S, Duran C, Thierer T, Ashton B, Meintjes P, Drummond A. 2012. Geneious Basic: an integrated and extendable desktop software platform for the organization and analysis of sequence data. Bioinformatics 28:1647–1649.

66. Hahn C, Bachmann L, Chevreux B. 2013. Reconstructing mitochondrial genomes directly from genomic next-generation sequencing reads--a baiting and iterative mapping approach. Nucleic Acids Res 41:e129.

67. Stamatakis A. 2014. RAxML version 8: a tool for phylogenetic analysis and post-analysis of large phylogenies. Bioinformatics 30:1312–1313.

68. Miller MA, Pfeiffer W, Schwartz T. 2010. Creating the CIPRES Science Gateway for inference of large phylogenetic trees, p. 1–8. In 2010 Gateway Computing Environments Workshop (GCE).

69. Buckley TR, Cordeiro M, Marshall DC, Simon C. 2006. Differentiating between hypotheses of lineage sorting and introgression in New Zealand alpine cicadas (*Maoricicada* Dugdale). Syst Biol 55:411–425.

70. Marshall DC, Slon K, Cooley JR, Hill KBR, Simon C. 2008. Steady Plio-Pleistocene diversification and a 2-million-year sympatry threshold in a New Zealand cicada radiation. Mol Phylogenet Evol 48:1054–1066.

71. Bator J, Marshall DC, Leston A, Cooley J, Simon C. 2021. Phylogeography of the endemic red-tailed cicadas of New Zealand (Hemiptera: Cicadidae: *Rhodopsalta*): molecular, morphological and bioacoustical confirmation of the existence of Hudson’s *Rhodopsalta microdora*. Zool J Linn Soc.

72. Weintraub PG, Beanland L. 2006. Insect Vectors of Phytoplasmas. Annu Rev Entomol 51:91–111.

73. Rakitov RA. 2002. Structure and Function of the Malpighian Tubules, and Related Behaviors in Juvenile Cicadas: Evidence of Homology with Spittlebugs (Hemiptera: Cicadoidea & Cercopoidea). Zoologischer Anzeiger - A Journal of Comparative Zoology 241:117–130.

74. Andersen PC, Brodbeck BV, Mizell RF. 1992. Feeding by the leafhopper, *Homalodisca coagulata*, in relation to xylem fluid chemistry and tension. J Insect Physiol 38:611–622.

75. Cheung WWK, Marshall AT. 1973. Water and ion regulation in cicadas in relation to xylem feeding. J Insect Physiol 19:1801–1816.

76. Williams KS, Simon C. 1995. The ecology, behavior, and evolution of periodical cicadas. Annu Rev Entomol 40:269–295.

77. Logan DP, Rowe CA, Maher BJ. 2014. Life history of chorus cicada, an endemic pest of kiwifruit (Cicadidae: Homoptera). N Z Entomol 37:96–106.

78. Buckley TR, Simon C. 2007. Evolutionary radiation of the cicada genus *Maoricicada* Dugdale (Hemiptera: Cicadoidea) and the origins of the New Zealand alpine biota. Biol J Linn Soc Lond 91:419–435.

79. Banker SE, Wade EJ, Simon C. 2017. The confounding effects of hybridization on phylogenetic estimation in the New Zealand cicada genus *Kikihia*. Mol Phylogenet Evol 116:172–181.

80. Groussin M, Mazel F, Sanders JG, Smillie CS, Lavergne S, Thuiller W, Alm EJ. 2017. Unraveling the processes shaping mammalian gut microbiomes over evolutionary time. Nat Commun 8:14319.

81. Sanders JG, Powell S, Kronauer DJC, Vasconcelos HL, Frederickson ME, Pierce NE. 2014. Stability and phylogenetic correlation in gut microbiota: lessons from ants and apes. Mol Ecol 23:1268–1283.

82. Wang J, Kalyan S, Steck N, Turner LM, Harr B, Künzel S, Vallier M, Häsler R, Franke A, Oberg H-H, Ibrahim SM, Grassl GA, Kabelitz D, Baines JF. 2015. Analysis of intestinal microbiota in hybrid house mice reveals evolutionary divergence in a vertebrate hologenome. Nat Commun 6:6440.

83. Brucker RM, Bordenstein SR. 2013. The Hologenomic Basis of Speciation : Science 466:667–669.

84. Chandler JA, Turelli M. 2014. Comment on “The hologenomic basis of speciation: gut bacteria cause hybrid lethality in the genus Nasonia.” Science.

85. Li Z, Wright A-DG, Si H, Wang X, Qian W, Zhang Z, Li G. 2016. Changes in the rumen microbiome and metabolites reveal the effect of host genetics on hybrid crosses. Environ Microbiol Rep 8:1016–1023.

86. Krell RK, Boyd EA, Nay JE, Park Y-L, Perring TM. 2007. Mechanical and Insect Transmission of *Xylella fastidiosa* to *Vitis vinifera*. Am J Enol Vitic 58:211–216.

87. Paião F, Meneguim AM, Casagrande EC, Lovato L, Leite RP. 2003. Levantamento de espécies de cigarras e transmissão de *Xylella fastidiosa* em cafeeiro.

88. Cummings NJ. 2009. Entomopathogenic fungi in New Zealand native forests: the genera Beauveria and Isaria. Doctor of Philosophy, University of Canterbury.

89. Price BW, Marshall DC, Barker NP, Simon C, Villet MH. 2019. Out of Africa? A dated molecular phylogeny of the cicada tribe Platypleurini Schmidt (Hemiptera: Cicadidae), with a focus on African genera and the genus *Platypleura* Amyot & Audinet-Serville. Syst Entomol 44:842–861.

90. Marshall DC, Moulds M, Hill KBR, Price BW, Wade EJ, Owen CL, Goemans G, Marathe K, Sarkar V, Cooley JR, Sanborn AF, Kunte K, Villet MH, Simon C. 2018. A molecular phylogeny of the cicadas (Hemiptera: Cicadidae) with a review of tribe and subfamily classification. Zootaxa 4424:1–64.

91. Hill KBR, Marshall DC, Marathe K, Moulds MS, Lee YJ, Pham T-H, Mohagan AB, Sarkar V, Price BW, Duffels JP, Schouten MA, de Boer AJ, Kunte K, Simon C. 2021. The molecular systematics and diversification of a taxonomically unstable group of Asian cicada tribes related to Cicadini Latreille, 1802 (Hemiptera : Cicadidae). Invertebr Syst 35:570–601.

92. McCutcheon JP, Boyd BM, Dale C. 2019. The Life of an Insect Endosymbiont from the Cradle to the Grave. Curr Biol 29:R485–R495.

